# Uncharted Waters: Projecting the European Emergence of *Naegleria fowleri*, Colloquially Known as the ‘Brain-Eating Amoeba’, Under Climate Warming

**DOI:** 10.64898/2026.09.02.748909

**Authors:** Volodymyr Tytar

## Abstract

**Background:** *Naegleria fowleri*, the causative agent of primary amoebic meningoencephalitis (PAM), is a thermophilic free-living amoeba historically endemic to warm freshwater environments, particularly in the southern United States. While human infections remain exceedingly rare, they are almost uniformly fatal. Global climate change is hypothesized to expand the geographic range of thermotolerant pathogens, potentially introducing them into previously unaffected regions.

**Objective:** This study aims to quantify the current and future habitat suitability for *N. fowleri* within the contiguous United States (US) and, critically, to project these models onto Europe—a continent traditionally considered low-risk—to identify regions of potential emergence under climate change scenarios.

**Methods:** We employed a modeling approach using the machine learning algorithm *Maxent*, implemented via the ‘flexsdm’ R package. We utilized 204 occurrence records from the contiguous US. We evaluated three environmental datasets (ENVIREM, Global Cloud Dynamics, Worldclim v.2), selecting Worldclim v.2 for its superior predictive performance. Future projections were generated for the 2041–2060 period under the Shared Socioeconomic Pathway (SSP) 2-4.5 (intermediate scenario). Variable importance was assessed using Shapley additive explanations (SHAP) to decouple the contribution of each Bioclimatic predictor and identify the key drivers of range expansion.

**Results:** The Worldclim v.2 dataset provided the most robust model. The model accurately identified the southeastern US as the core current habitat, consistent with historical PAM case distribution. Crucially, when projected onto Europe, the model reveals extensive regions of currently low suitability that transition to moderate or high suitability by 2050, particularly across the Mediterranean basin, parts of Central Europe, and the Atlantic coastal fringes. SHAP analysis identified temperature-related variables —specifically Annual Mean Temperature, Mean Temperature of the Coldest Quarter, and Mean Diurnal Range—as the most influential predictors, followed closely by precipitation variables (Precipitation of Driest Quarter/Month and Precipitation Seasonality). Under the SSP2-4.5 scenario, the model projects a distinct northward and eastward expansion of suitable habitat, with Ukraine emerging as a region of particular concern, especially in light of recent anthropogenic hydrological changes to the Dnipro River basin.

**Conclusion:** Our findings support the hypothesis that global warming will facilitate the geographic spread of *N. fowleri*, carrying this pathogen into uncharted European waters. The projections for Europe—and for Ukraine in particular—suggest a non-negligible risk of future autochthonous PAM cases. These results warrant the proactive inclusion of this pathogen in public health surveillance programs and climate change adaptation strategies across the continent.

## 1. Introduction

*Naegleria fowleri* is a free-living, thermophilic amoeboflagellate found globally in soil and warm freshwater bodies (e.g., lakes, rivers, hot springs, and poorly chlorinated swimming pools)(DeFelice, 2026). Human infection occurs when water containing the amoeba is forcefully introduced into the nasal cavity, allowing the trophozoite to migrate along the olfactory nerve to the brain. This results in primary amoebic meningoencephalitis (PAM), a rapidly progressive and devastating disease with a case fatality rate exceeding 97% (Capewell et al., 2015).

Historically, the majority of PAM cases have been reported from the southern-tier states of the US, Australia, and tropical regions. However, the protozoan’s optimal growth temperature ranges from 30°C to 46°C, making it acutely sensitive to environmental thermal regimes (Maciver et al., 2020). Climate change, characterized by rising average global temperatures and increased frequency of heatwaves, is expected to alter the distribution of thermophilic pathogens. Warmer air and water temperatures may extend the transmission season and permit the amoeba to establish itself in previously unsuitable, cooler latitudes.

Europe has generally been considered a low-risk zone for *N. fowleri*, with only sporadic imported or isolated cases reported (Indolfi et al., 2024). However, recent summer heatwaves across the continent have raised concerns about the potential endemic establishment of the pathogen. Predictive niche modeling offers a powerful tool to proactively assess this threat. Machine learning algorithms, such as, for instance, Maximum Entropy (*Maxent*). This machine learning framework has successfully been used in ecological niche modeling (ENM) to predict the geographic distribution and habitat suitability of pathogens and free-living amoebae (FLA). Because it relies purely on “presence-only” data (locations where the organisms have been detected) and environmental layers, *Maxent* is ideal for mapping cryptic microscopic organisms where definitive “absence” data is nearly impossible to confirm (Stahl et al., 2025; Kavur et al., 2026).

The claim that *Maxent* is a powerful and widely used tool for predicting pathogen distributions is strongly supported by a number of studies (for example, Khalaf et al., 2024; Alqahtani et al., 2025; Khalaf et al., 2025; Zheng et al., 2026). They collectively demonstrate that *Maxent* is the standard and preferred method for this type of ecological niche modeling (ENM), especially when working with presence-only data for pathogens.

In this study, we leverage the comprehensive occurrence data available for the contiguous United States — where the pathogen is relatively well-documented — to build robust habitat suitability models. We then project these models onto Europe to identify habitable regions that could become suitable under future climate scenarios, making a special emphasis on Ukraine. We utilize the flexible species distribution modeling framework (‘flexsdm’)(Velazco et al., 2022) and enhance model interpretability using SHAP values (Wang et al., 2024) to identify the key climatic drivers of this potential range expansion.

## 2. Materials and Methods

### 2.1. Occurrence Data

We used published occurrence records for *N. fowleri* from environmental samples (water/soil) and confirmed PAM cases across the contiguous United States (Stahl et al., 2025). 204 occurrence records for the contiguous US were utilized. After cleaning the data in SAGA GIS (Conrad et al., 2015), a final set of 96 unique (i.e., non-duplicate) occurrence points was resorted for model calibration. All geographic coordinates (in decimal degrees) were verified and standardized using Google Maps (https://www.google.com/maps). Subsequently, the data were transformed into comma-separated values (CSV) format in preparation for analysis.

### 2.2. Environmental Predictors

We tested three distinct environmental datasets to capture the Bioclimatic and hydrologic niche of the species.

1. *ENVIREM (v2*.*0)*, focuses on water balance and thermal extremes (Title, Bemmels, 2018).
2. *EarthEnv Global 1-km Cloud Frequency Version 1*, incorporates variables related to solar radiation and cloud cover (Wilson, Jetz, 2016). Cloud cover can influence numerous important ecological processes including reproduction, growth, and survival, cloud cover dynamics may provide key information for delineating a variety of habitat types and predicting species distributions.
3. *Worldclim (v2*.*1)* provides 19 standard Bioclimatic variables derived from monthly temperature and precipitation values (Fick, Hijmans, 2017). These variables represent annual trends, seasonality, and extreme or limiting environmental factors commonly used in ecological and species distribution modeling.

### 2.3. Modeling Framework

Within the flexible species distribution modeling framework for projecting the potential geographic distribution of N. fowleri spatial block partitioning was used to generate pseudo-absence and background points. Filtering the occurrence data was used to reduce sample bias by randomly removing points where they were dense (oversampling) in the environmental and geographical spaces. The ‘flexsdm’ package offers a wide range of modeling options. Here, we tested out Maximum Entropy (*Maxent*) (Phillips et al., 2006), one of the most popular SDM modelling methods. *Maxent* can construct simple to highly complex, nonlinear species–environment relationships using various transformations of variables termed features and representedby a number of feature classes (FC) of which we tested linear (L), quadratic (Q) and product (P), and their combinations (LQ, LP, QP, LQP). To reduce overfitting, *Maxent* uses a regularization procedure to balance model fit with complexity, by penalising models based on the magnitude of their coefficients. Tuned models were built using regularization multiplier values ranging from 0 to 4 with increments of 0.5. The model’s predictive accuracy was measured using the widely recognised AUC statistic. AUC scores range from 0 to 1, with values closer to 1 reflecting strong discriminatory power in distinguishing habitat suitability (Fielding, Bell, 1997) and the true skill statistic (TSS) where the value of >0.4 is considered good, with the range of 0.4–0.8 indicating ‘good’ performance; a score of >0.8 is considered excellent (Allouche et al., 2006). But whereas AUC remains a controversial criterion (Lobo, 2008), for greater confidence we employed the continuous Boyce index, CBI (Boyce et al., 2002), one of the most reliable presence-only evaluation metrics, and also provided by the ‘flexsdm’ package. It varies between −1 and +1. Positive values indicate a model that presents predictions that are consistent with the distribution of presences in the evaluation dataset, values close to zero mean that the model is not different from a random model (Hirzel et al., 2006).

All projection outputs were generated in GeoTIFF format and subsequently imported into SAGA GIS for cartographic visualization to facilitate comparative analysis across regions and time periods. For Ukraine we calculated the areal extent of suitable habitat (defined as areas exceeding the maximum training sensitivity plus specificity threshold) to quantify the magnitude of projected range expansion. Statistical data was analysed using the PAST software package (Hammer et al., 2001) and the R environment (https://www.r-project.org).

### 2.4. Conditioning factors

Commonly used approaches recommend removing correlated predictor variables before modeling to avoid multicollinearity, which is considered to affect model projections (Zhao et al., 2022), and there are several statistical packages offering functions that reduce collinearity in predictors. However, we avoided removing correlated predictor variables because the benefits of using all available variables may outweigh the drawbacks of collinearity. Latest research indicates that modeling with correlated climate variables increases accuracy of predictions (Hanberry, 2023). Moreover, complex models such as *Maxent* take advantage of existing collinearity in finding the best set of parameters (De Marco, Nóbrega, 2018).

#### 2.4.1. Variable Importance with SHAP

To further explore the impact of the considered above environmental factors, we employed a SHAP framework from XAI (i.e., eXplainable artificial intelligence) to rank and uncover the most influential drivers (Lundberg et al., 2018; Farooq et al., 2022). With a SHAP approach there is no need to consider only uncorrelated environmental drivers (Nikraftar et al., 2025). SHAP (SHapley Additive exPlanations) is a unified framework in explainable artificial intelligence used to interpret the output of any machine learning model by assigning each feature an importance value for a particular prediction. Usung the shapviz: SHAP Visualizations R package version 0.10.4 (https://github.com/ModelOriented/shapviz), we post-processed the best model results with SHAP by comparing what a model predicts with and without the predictor for all possible combinations of predictors at every single observation. The predictors are then ranked according to their contribution for each observation and averaged across observations. Complementing the global importance rankings, SHAP dependence plots were further generated to visualize the marginal effect of each top-ranked predictor on predicted suitability, revealing nonlinear thresholds and interaction effects that are not captured by coefficient-based metrics alone. The application of SHAP for understanding the influence of environmental factors on species distribution is now being investigated more widely (e.g., Buebos-Esteve, Dagamac, 2025).

### 2.5. Model Projections

Following model preparation, we generated spatial projections of habitat suitability for *N. fowleri* under both current and future climate conditions. These projections were implemented using the calibrated *Maxent* models within the ‘flexsdm’ framework, with environmental layers resampled to a common spatial resolution and clipped to the respective geographical extents of interest.

Current climate projections were produced by applying the final models to the baseline environmental layers mentioned above. These projections were first generated for the contiguous United States, serving both as a means of model validation (by comparing predicted suitability against known occurrence records and historical PAM case distribution) and as a baseline reference for subsequent transcontinental extrapolation. The resulting habitat suitability maps express the relative probability of species presence on a continuous scale from 0 (lowest suitability) to 1 (highest suitability), allowing for the identification of core endemic areas and ecological thresholds.

Future climate projections were developed using climate forcing data for the 2041–2060 period (hereafter referred to as the 2050s), derived from the Coupled Model Intercomparison Project Phase 6 (CMIP6) under the Shared Socioeconomic Pathway SSP2-4.5. This scenario represents a “middle-ofthe-road” trajectory, projecting moderate greenhouse gas emissions and intermediate levels of socioeconomic challenges to adaptation and mitigation. Under SSP2-4.5, global mean temperatures are expected to increase by approximately 2.0–2.5°C above pre-industrial levels by the end of the century, making this scenario a plausible and policy-relevant benchmark for assessing medium-term climatic shifts. The same ensemble model was applied to these future layers to project habitat suitability across both the contiguous United States and, more critically, the European continent. This dual projection approach serves two purposes: (i) it allows for a direct comparison of predicted range dynamics within the model’s training domain, and (ii) it enables the identification of emerging risk areas in Europe—a region where *N. fowleri* occurrence data remain sparse but where climate trends suggest increasing thermal suitability.

## 3. Results

Based on model performance, Worldclim v.2.1 unambiguously was selected as the definitive environmental dataset due to its superior explanatory power and relevance to the species’ known thermal tolerances (CBI=0.90+/-0.07SD).

### 3.1. Model Performance and Variable Contributions

The ensemble *Maxent* model (FC=L, regularization multiplier=1), constructed by averaging the top-performing tuned models (based on AUC and TSS scores), demonstrated robust predictive performance across all evaluation metrics. The mean Area Under the Receiver Operating Characteristic Curve (AUC) reached 0.85+/-0.04SD, indicating excellent discriminatory capacity in distinguishing suitable from unsuitable habitats. The True Skill Statistic (TSS) yielded a value of 0.71+/-0.07, substantially exceeding the conventional threshold of 0.4 considered indicative of good model performance. Furthermore, the continuous Boyce index (CBI) registered 0.96+/-0.01, confirming that the predicted suitability values were consistently and positively correlated with the distribution of presence records in the evaluation dataset. Collectively, these metrics attest to the model’s strong predictive reliability and its suitability for current-climate transcontinental projection.

#### 3.1.1. Current Habitat Suitability: US and European Baseline

This baseline serves a dual purpose: it validates the model’s predictive capacity by comparing outputs against known endemic areas, and it provides a reference against which future climate-driven changes can be quantified.

The baseline suitability map for the contiguous US accurately reproduced the species’ historically documented endemic range, demonstrating the model’s strong calibration and its ability to capture the macroclimatic determinants of *N. fowleri* distribution in this part of the world. High-suitability areas (probability > 0.7) were concentrated across the southeastern states, including Florida, Texas, Louisiana, Georgia, and extending into adjacent portions of Alabama, Mississippi, and South Carolina. Additional pockets of high suitability were identified in the southwestern United States, particularly in parts of Arizona and New Mexico, where warm thermal regimes and the presence of suitable freshwater habitats support amoebal populations. This spatial pattern aligns closely with the geographic distribution of historically reported primary amoebic meningoencephalitis (PAM) cases, which have predominantly occurred in the southern-tier states during summer months when water temperatures peak. The congruence between predicted suitability and observed occurrences provides strong empirical support for the model’s ecological validity and its utility for extrapolation to other geographical regions.

When projected onto the European continent under current climatic conditions, the model revealed a different landscape of suitability. The vast majority of Europe, particularly the northern and eastern regions, was classified as low suitability (probability < 0.3), reflecting the prevailing cooler thermal regimes that currently constrain the species’ distribution. This includes Scandinavia, the Baltic states, Belarus, most of the European part of Russia, and the northern reaches of Central Europe, where cold winter temperatures and moderate summer warmth fall below the physiological thresholds required for firm *N. fowleri* establishment. Similarly, much of Eastern Europe—including Poland, Ukraine, Slovakia, Hungary, and the Czech Republic—exhibited low to marginal suitability under baseline conditions.

Pockets of moderate suitability (probability 0.3–0.6) were identified in the southern Mediterranean basin, where warmer climatic conditions create more favorable thermal environments. These include the southern Iberian Peninsula (particularly the Guadalquivir Valley in Spain and the Algarve region of Portugal), southern Italy (including Sicily and Calabria), and the Greek mainland and islands. In these regions, the combination of warm annual temperatures, mild winters, and sufficient precipitation supports aquatic habitats that may already be marginally suitable for *N. fowleri* survival, though the species has rarely been documented in these areas to date. The absence of reported cases likely reflects a combination of limited surveillance, lower population densities in high-risk areas, and the fact that thermal conditions currently approach but do not consistently exceed the thresholds required for sustained transmission.

**Fig 1.**
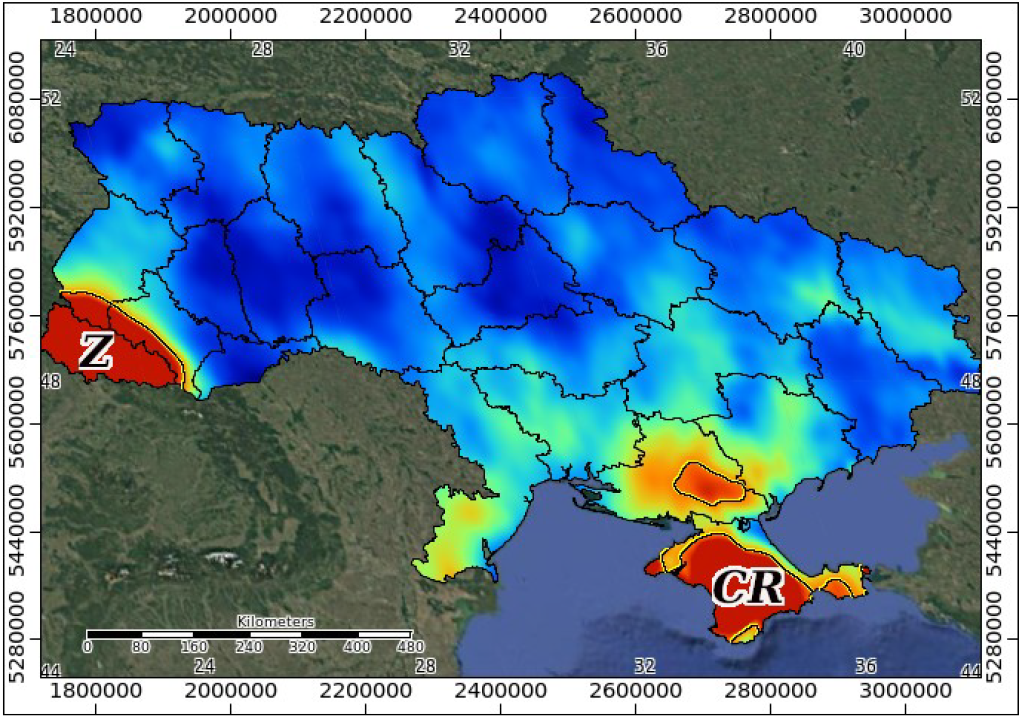
Projected habitat suitability (HS) map for *Naegleria fowleri* in Ukraine based on the current time period (*Worldclim (v2*.*1)*; colours show potential HS ranging from high (red) to low (blue); the yellow/black-coloured contour line represents the maximum training sensitivity plus specificity threshold; Z - Zakarpatska Oblast, CR- Crimea. Coordinate reference system: Mollweide projection.

Within the broader European context, Ukraine exhibits a predominantly low-suitability profile under current climate conditions (projected mean habitat suitability=0.25). However, the model identified two discrete regions of elevated suitability that warrant specific attention. The first is Transcarpathia (Zakarpattia Oblast) in the far western part of the country, where the region’s relatively mild winters— attributable to its proximity to the Pannonian Plain and the moderating influence of the Carpathian Mountains—result in marginally higher suitability compared to surrounding territories. The second is the Crimean Peninsula (currently occupied by Russia), where the warmer Mediterranean-influenced climate of the southern coast and the Black Sea basin creates conditions that are more conducive to *N. fowleri* presence. These localized pockets of moderate suitability underscore the heterogeneous nature of current climate risk within Ukraine and suggest that, while the country as a whole remains at low baseline risk, certain subregions may already possess environmental conditions approaching the species’ thermal and hydrological requirements.

In summary, the baseline projections reveal a clear latitudinal and climatic gradient in current *N. fowleri* suitability across both North America and Europe. The southeastern US represents the core endemic range, while Europe is largely characterized by unsuitable or marginal conditions, with the exception of Mediterranean coastal fringes and limited microclimatic zones in southern and western Ukraine.

#### 3.1.2. Future Projections for Europe

Projecting the ensemble *Maxent* model (FC=LQ, regularization multiplier=2) onto future climate conditions for the 2041–2060 period under the Shared Socioeconomic Pathway SSP2-4.5 (AUC=0.85+/-0.14SD, TSS=0.73+/-0.18SD, CBI=0.95+/-0.01SD) revealed a pronounced and spatially extensive transformation of *N. fowleri* habitat suitability across the European continent. The contrast between current and future projections is striking, with the model predicting a substantial northward and eastward expansion of suitable environments, driven primarily by rising temperatures—particularly during winter months—and shifts in precipitation patterns that collectively relax the climatic constraints currently limiting the species’ distribution.

##### Mediterranean Basin and Southern Europe

Under the warming scenario, the Mediterranean basin emerges as the epicenter of future habitat expansion. High-suitability zones (probability > 0.7) are projected to develop across the entire Mediterranean coastline, encompassing the southern Iberian Peninsula, the Balearic Islands, coastal and insular Italy (including Sicily and Sardinia), the Greek mainland and Aegean islands, the Mediterranean coast of France, and the Adriatic and Ionian coasts of the Balkans. These regions are characterized by projected increases in annual mean temperature of approximately 2–3°C, coupled with milder winter minima that consistently exceed the critical thresholds for cyst survival. The intensification of suitability in these areas suggests that, by midcentury, much of southern Europe will possess environmental conditions comparable to those currently found in the southeastern United States—the species’ historical endemic core.

##### Northward Expansion into Central and Western Europe

The model predicts a significant northward penetration of suitable habitats beyond the Mediterranean basin, with high-suitability corridors extending into major river valleys and lowland regions that combine favorable thermal regimes with hydrological stability. The Po Valley in northern Italy is projected to become highly suitable, reflecting the region’s warm summers, mild winters, and extensive network of freshwater bodies that provide stable aquatic habitats. Similarly, the Rhône Valley in southeastern France and the Rhine Basin in Germany and the Benelux countries are identified as zones of emerging high suitability. These riverine corridors are particularly concerning from a public health perspective, as they are densely populated regions with extensive recreational water use and complex water infrastructure. The Rhine Basin, in particular, represents a significant northward leap in suitability, indicating that central Europe may no longer serve as a climatic barrier to *N. fowleri* establishment by mid-century.

##### Temperate and Atlantic Regions

Temperate regions of Western and Northern Europe, including the United Kingdom, Ireland, the Benelux countries, and France, are projected to shift from low suitability under current conditions to moderate suitability (probability 0.3–0.6) by the 2050s. This transition is primarily attributable to warmer winter temperatures that eliminate or reduce the cold-season bottlenecks currently preventing population persistence. While moderate suitability does not imply the same level of risk as high-suitability zones, it nevertheless suggests that environmental conditions may become permissive enough to support sporadic amoebal presence, particularly during summer heatwaves when water temperatures can locally exceed growth thresholds. The model further indicates that southern Scandinavia—including the southern tips of Sweden and Norway—may shift from very low to low-moderate suitability, representing the northernmost limit of projected expansion. This poleward shift underscores the pervasive influence of climate warming on the redistribution of thermophilic organisms, even in regions historically considered climatically inhospitable.

**Fig 2.**
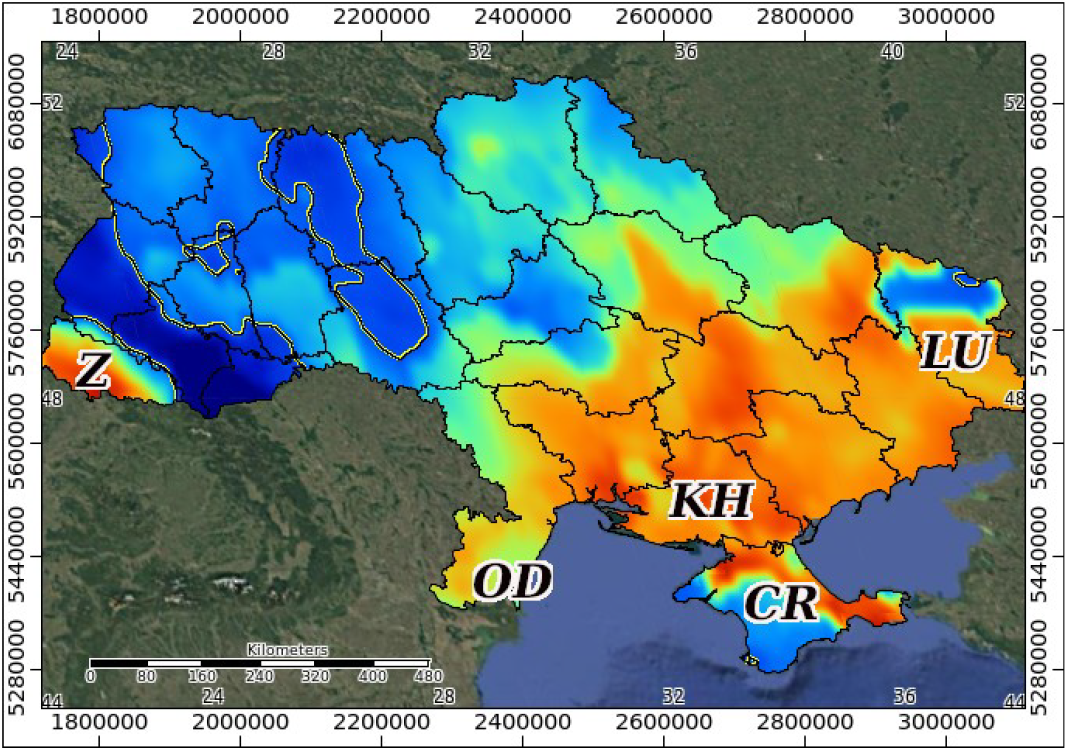
Projected habitat suitability (HS) map for *Naegleria fowleri* in Ukraine based on the future time period 2041–2060 (*Worldclim (v2*.*1)*; colours show potential HS ranging from high (red) to low (blue); the yellow/black-coloured contour line represents the maximum training sensitivity plus specificity threshold; Z - Zakarpatska, OD - Odeska, KH - Khersonska, LU - Luhanska oblasts, CR- Crimea. Coordinate reference system: Mollweide projection.

##### Eastern Europe and Ukraine

Eastern Europe, including the Baltic states, Poland, the Czech Republic, Slovakia, Hungary, and the Balkan interior, is projected to experience a general increase in suitability, with many areas transitioning from low to moderate suitability under the SSP2-4.5 scenario. However, the most notable changes within this region are projected for Ukraine, where the model predicts a significant eastward and southward expansion of suitable habitats and double rise in habitat suitability (mean habitat suitability=0.54). Under current conditions, suitable areas in Ukraine are largely confined to Transcarpathia in the west and the Crimean Peninsula in the south. By the 2050s, the model projects the emergence of a contiguous belt of moderate to high suitability stretching across the southern and southeastern regions of the country, extending from Odesa Oblast in the southwest along the Black Sea coast and inland through the steppe and forest-steppe zones to Luhansk Oblast in the east. This expansion encompasses the Dnipro River corridor and its associated reservoirs and irrigation canals, which may provide the stable freshwater environments required for amoebal colonization. The projected increase in suitability across southern and eastern Ukraine reflects the combined effects of warming winter and summer temperatures, reduced frost frequency, and the region’s relatively low precipitation seasonality, which supports persistent aquatic habitats. Given Ukraine’s extensive agricultural and industrial water infrastructure, including the Kakhovka and Dnipro reservoirs (particularly the critically damaged first one: Shumilova et al., 2025; Vyshnevskyi et al., 2023), this projected expansion warrants particular attention from public health and water management authorities. *General Patterns and Ecological Implications*. Across the European continent, the future projections reveal a consistent pattern of range expansion that is both latitudinal (northward) and longitudinal (eastward). The model indicates that the zone of high suitability will effectively extend from the Mediterranean coastline northward into central Europe, while moderate suitability will expand into regions previously considered climatically unsuitable. Notably, the greatest relative increases in suitability are projected for areas currently at the cold edge of the species’ environmental envelope, where small absolute increases in winter temperature produce disproportionately large gains in predicted suitability. This nonlinear response is consistent with the threshold-type relationships identified below in the SHAP analysis for temperature variables—particularly the Mean Temperature of the Coldest Quarter (Bio11), which exhibited a steep increase in suitability once winter temperatures exceed approximately 10–12°C. The implication is that even moderate climate warming scenarios, such as SSP2-4.5, can trigger substantial ecological shifts in thermophilic species distributions, highlighting the potential for abrupt rather than gradual changes in pathogen emergence risk.

### 3.2. SHAP analysis of variable importance

To move beyond conventional variable importance rankings and gain deeper mechanistic insight into the ecological drivers of *N. fowleri* distribution, we applied SHAP (SHapley Additive exPlanations) analysis to the final ensemble model. Unlike traditional permutation-based importance measures, SHAP provides a unified and theoretically grounded framework for quantifying the marginal contribution of each predictor to the model’s predictions across all individual observations. This approach allows for the decomposition of the model output into additive feature contributions, revealing not only the relative ranking of variables but also the direction and shape of their relationships with habitat suitability. The SHAP analysis yielded a clear hierarchy of variable importance, with six Bioclimatic factors emerging as the primary determinants of *N. fowleri* habitat suitability. These are discussed below in descending order of influence.

1. *Annual Mean Temperature* (Bio1) ranked as the single most influential predictor, with SHAP values demonstrating a strongly positive and approximately exponential relationship with habitat suitability. This finding is ecologically consistent with the known thermophilic physiology of *N. fowleri*, whose trophozoites exhibit optimal growth between 30°C and 46°C and fail to proliferate below approximately 25°C (for example, Yoder et al., 2010). The model indicates that regions with annual mean temperatures exceeding approximately 15–17°C become increasingly suitable, with suitability rising sharply above the 20°C threshold. This variable effectively captures the broad latitudinal and continental thermal gradients that constrain the species’ fundamental niche, explaining why historically endemic regions are concentrated in subtropical and warm-temperate zones.
2. *Mean Temperature of the Coldest Quarter* (Bio11) emerged as the second most important predictor, underscoring the critical role of winter thermal regimes in determining the species’ capacity for perennial establishment. SHAP analysis revealed a pronounced threshold effect: suitability remains negligible where the mean temperature of the coldest quarter falls below approximately 5°C, increases gradually between 5°C and 10°C, and rises steeply above 12°C. This pattern is Biologically meaningful, as it reflects the temperature-dependent survival of cysts—the dormant stage that enables the amoeba to overwinter in sediments and re-initiate the life cycle when water temperatures rise in spring and summer. Mild winters, therefore, represent a prerequisite for the maintenance of stable, self-sustaining populations, and the projected warming of winter temperatures under climate change is likely to be a primary driver of range expansion into higher latitudes. 3&4. *Precipitation of the Driest Quarter* (Bio17) and *Precipitation of the Driest Month* (Bio14) ranked third and fourth in importance, respectively, and both exhibited negative correlations with habitat suitability. The SHAP dependence plots for these variables showed that suitability decreases markedly when precipitation during the driest quarter falls below approximately 50 mm, with a further decline observed when the driest month receives less than 15–20 mm. This relationship likely reflects the hydrological requirements of the species: *N. fowleri* inhabits freshwater bodies such as lakes, ponds, rivers, and thermally polluted waters, all of which depend on sufficient and consistent water availability. Prolonged dry periods may reduce the extent, depth, and thermal stability of aquatic habitats, leading to desiccation, increased salinity, or elevated temperatures beyond the species’ upper tolerance limit. Conversely, areas with adequate dry-season precipitation are more likely to maintain the stable water bodies that support amoebal populations. The slightly higher importance of Bio17 compared to Bio14 suggests that the cumulative water deficit over the entire driest quarter is a more critical filter than the minimum monthly value alone.
3. *Mean Diurnal Range* (Bio2)—defined as the mean of monthly differences between daily maximum and minimum temperatures—ranked fifth in the SHAP hierarchy. This variable exhibited a moderate and predominantly negative influence on suitability, with SHAP values indicating a gradual decline in suitability as the diurnal temperature range increases beyond approximately 10–11°C. A narrower diurnal range is characteristic of regions with higher atmospheric moisture, greater cloud cover, or proximity to large water bodies, all of which buffer against rapid daily temperature fluctuations. Such conditions may favor *N. fowleri* by reducing thermal stress on trophozoites and promoting more stable thermal environments in shallow freshwater habitats. In contrast, regions with large diurnal swings (typical of continental interiors) may experience temperature regimes that frequently exceed the species’ upper thermal tolerance during the day or drop below growth thresholds at night, thereby limiting population persistence.
4. *Precipitation Seasonality* (Bio15), expressed as the coefficient of variation in monthly precipitation totals, was the sixth most influential predictor. Its relationship with suitability was more complex, with SHAP values suggesting an optimal range rather than a simple monotonic trend. Suitability peaked at intermediate levels of precipitation seasonality (coefficient of variation approximately 40–60%) and declined under both highly seasonal (strong wet-dry contrasts) and very aseasonal (extremely uniform) precipitation regimes. This pattern likely reflects a trade-off: moderate seasonality ensures sufficient water availability during the growing season while preventing the extremes of prolonged drought or excessive dilution and flushing that could destabilize aquatic habitats. Regions with pronounced dry seasons may experience habitat contraction, while regions with overly uniform precipitation may lack the thermal stratification or nutrient dynamics that favor amoebal proliferation. In summary, the SHAP analysis robustly identifies temperature as the primary climatic filter governing *N. fowleri* distribution, with annual means and winter minima exerting the strongest constraints. Precipitation variables act as secondary but nonetheless essential modifiers, regulating the hydrological stability of freshwater habitats. This hierarchy of variable importance aligns closely with the known eco-physiological characteristics of the species and provides a mechanistic foundation for interpreting the projected range dynamics under future climate scenarios. Notably, the dominance of temperature-related predictors—particularly those associated with cold-season conditions—corroborates the hypothesis that climate warming will be the principal driver of poleward range expansion, as previously hypothesized in the literature (Maciver et al., 2020; Stahl et al., 2025).

**Fig 3.**
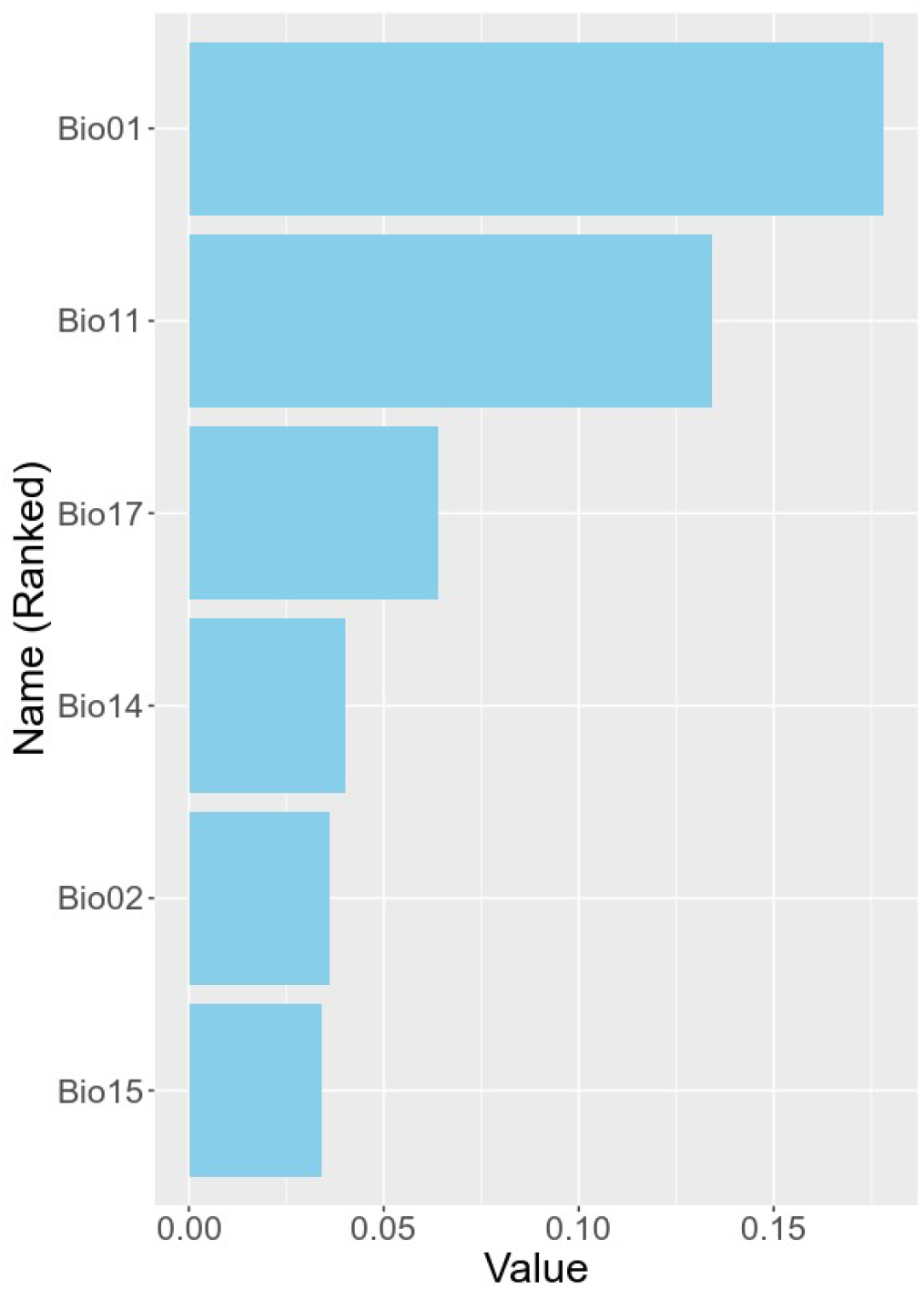
Summary bar-plots of the average absolute SHAP values of the six top contributing to the Maxent model variables. The y-axis represents variables used in the study: *Annual Mean Temperature* (Bio01) [0.178], *Mean Temperature of the Coldest Quarter* (Bio11) [0.134], *Precipitation of the Driest Quarter* (Bio17) and *Precipitation of the Driest Month* (Bio14) [0.064 & 0.040], *Mean Diurnal Range* (Bio02) [0.036], *Precipitation Seasonality* (Bio15) [0.034]. The x-axis represents the corresponding SHAP value (in []).

## 4. Discussion

This study provides the first quantitative, spatially explicit projection of *N. fowleri* habitat suitability in Europe under a climate change scenario. The use of the contiguous US as a model training ground is justified by the relative abundance of occurrence data, and the high performance of our models validates this transatlantic projection. Our findings corroborate the hypothesis that climate warming will facilitate the geographic spread of thermotolerant pathogens, with Europe—and particularly the Mediterranean basin and parts of Eastern Europe—emerging as regions of increasing concern.

### 4.1. Climatic Drivers of Expansion

The dominance of temperature variables in the SHAP analysis confirms that *N. fowleri* is fundamentally a thermophilic organism. The critical finding is the shift in the “Mean Temperature of the Coldest Quarter.” As winters in Europe warm (projected to increase by 2–3°C under SSP2-4.5), previously lethal cold thresholds for cyst survival will be surpassed, allowing the amoeba to establish perennial populations. The role of precipitation—specifically the significance of the driest quarter— suggests that water availability and hydrological stability are secondary filters. Warmer winters combined with stable summer water levels are the ideal conditions for the amoeba to thrive. This mechanistic understanding, derived from SHAP analysis, provides a robust foundation for interpreting the projected range dynamics and for identifying regions most vulnerable to future emergence.

### 4.2. The Kakhovka Reservoir Transformation as a Case Study in Emerging Risk

Our projections for Ukraine indicate a significant expansion of suitable habitat into southern and eastern regions by mid-century. This projected risk is substantially amplified by the profound hydrological transformation resulting from the destruction of the Kakhovka Dam in June 2023. The former Kakhovske Reservoir, once a large, deep water body with considerable thermal inertia and a stable temperature profile, covered a total area of 2,155 km^2^ in the Kherson, Zaporizhzhia, and Dnipropetrovsk oblasts of Ukraine, has been drained and replaced by a fragmented network of shallow river branches, residual lakes, and isolated pools.

Scientific studies have documented this dramatic change, demonstrating that the newly formed water bodies are fundamentally different from the original reservoir. Research by V. Vyshnevskyi and coauthors (2023) shows that “the former Kakhovske reservoir became a network of river branches and lakes that cannot be compared with the former reservoir”. This fragmentation has profound hydrological and ecological implications: the reduced water volume and increased surface-area-to-volume ratio mean that these new, shallow water bodies are much more sensitive to atmospheric heating. Prior to the dam’s destruction, studies had already documented a warming trend in Dnipro River reservoirs, with summer water temperatures increasing by 0.74°C per decade between 1977 and 2020 . The post-destruction landscape will likely accelerate this warming, as the shallow, isolated pools heat up more rapidly and reach higher temperatures during summer heatwaves.

The ecological consequences of this transformation extend beyond temperature. The newly exposed sediments of the former reservoir bed contain a legacy of pollutants—including heavy metals, petroleum by-products, and other toxic contaminants—accumulated over decades of industrial activity. As noted by Shumilova et al. (2025), this contamination “poses a largely overlooked long-term threat to freshwater, estuarine, and marine ecosystems”. Furthermore, the draining of the reservoir has caused an ecological disaster for approximately 40 species of fish, with the remaining water surface reduced to about 19% of its original area within two months of the disaster.

The relevance of these hydrological changes to *N. fowleri* risk is direct and concerning. The newly formed shallow water bodies—warmer, slower-flowing, and more nutrient-rich due to sediment disturbance—create conditions that align precisely with the ecological niche of this thermophilic amoeba. As our model projects increasing suitability for *N. fowleri* in southern Ukraine under climate warming, the actual risk may be further elevated by these local hydrological transformations. This represents a potent example of how climate-driven range expansion can interact with anthropogenic environmental change to create novel hotspots of pathogen risk. Public health and water management authorities in the region should prioritize the monitoring of these newly formed water bodies, particularly during summer months when water temperatures peak and recreational water use is highest.

### 4.3. Public Health Implications for Europe

The projected expansion does not imply an imminent epidemic; PAM remains a rare disease. However, the identification of future high-suitability zones—such as the Mediterranean coast and, increasingly, parts of Eastern Europe—has several critical implications:

1. *Clinical Awareness*. Physicians in these regions should become aware of PAM as a differential diagnosis in patients presenting with acute meningitis and a history of freshwater swimming. Early recognition is crucial, as prompt initiation of investigational therapies may improve outcomes, though the case fatality rate remains exceedingly high.
2. *Recreational Risk*. Public health advisories regarding warm, stagnant freshwater bodies during summer heatwaves may need to be issued. This is particularly relevant for the newly formed water bodies in the former Kakhovka Reservoir area, where warmer conditions and increased public recreation could elevate exposure risk.
3. *Water Infrastructure*. Drinking water and treated recreational water systems (e.g., splash pads, swimming pools) in warming regions may require enhanced chlorine monitoring to prevent amoebal colonization. The use of untreated freshwater for nasal irrigation or other purposes should be strongly discouraged.

### 4.4. Limitations

This study has limitations inherent to correlative niche modeling. We cannot account for adaptive evolution of the amoeba or its microBiome. Furthermore, the occurrence data from the US may not capture the full physiological niche of the species, which could differ slightly in European strains. The use of a single climate scenario (SSP2-4.5) provides a moderate outlook; more extreme scenarios would likely accelerate the projected spread. Additionally, our projections do not explicitly account for the hydrological changes described above, meaning that local risk in areas like the former Kakhovka Reservoir may be underestimated. Future work should integrate high-resolution hydrological and water temperature data to refine these projections at the local scale.

## 5. Conclusion

Our modeling approach, utilizing *Maxent* and SHAP analysis, robustly demonstrates that climate warming will significantly expand the environmental envelope of *N. fowleri*. The projections for Europe indicate a clear trend: a northward and eastward expansion of suitable habitats by the mid-21st century. While PAM will likely remain a rare event, the expanding spatial risk necessitates proactive surveillance and preparation. This study underscores the profound impact of climate change on the distribution of infectious pathogens and advocates for the integration of disease ecology models into public health policy.

